# The large terminase DNA packaging motor grips DNA with its ATPase domain for cleavage by the flexible nuclease domain

**DOI:** 10.1101/080440

**Authors:** Brendan J. Hilbert, Janelle A. Hayes, Nicholas P. Stone, Rui-Gang Xu, Brian A. Kelch

## Abstract

Many viruses use a powerful terminase motor to pump their genome inside an empty procapsid shell during virus maturation. The large terminase (TerL) protein contains both enzymatic activities necessary for packaging in such viruses: the ATPase that powers DNA translocation and an endonuclease that cleaves the concatemeric genome both at initiation and completion of genome packaging. However, how TerL binds DNA during translocation and cleavage is still mysterious. Here we investigate DNA binding and cleavage using TerL from the thermophilic phage P74-26. We report the structure of the P74-26 TerL nuclease domain, which allows us to model DNA binding in the nuclease active site. We screened a large panel of TerL variants for defects in binding and DNA cleavage, revealing that the ATPase domain is the primary site for DNA binding, and is required for nucleolysis. The nuclease domain is dispensable for DNA binding but residues lining the active site guide DNA for cleavage. Kinetic analysis of nucleolysis suggests flexible tethering of the nuclease domains during DNA cleavage. We propose that interactions with the procapsid shell during DNA translocation conformationally restrict the nuclease domain, inhibiting cleavage; TerL release from the procapsid upon completion of packaging unlocks the nuclease domains to cleave DNA.

## Introduction

Most double-stranded DNA viruses package their genomes using an ATP-dependent motor to pump DNA into an empty capsid protein shell. As DNA fills the shell, internal pressure builds due to confinement of the highly charged DNA. Therefore, these motors have evolved to become some of the most powerful bio-motors known (1,2). For this reason, there is much interest in engineering packaging motors for delivery of nucleic acid therapeutics and as functionalized nano-materials. Moreover, genome packaging motors from herpes viruses are the targets of various FDA-approved antiviral drugs (3–9).

There are two distinct families of packaging motors for membrane-free dsDNA viruses: the terminase family and the Phi29-family motors (10). Here we focus on the more common terminase packaging apparatus, which has been studied in many viral systems (11–18). Terminase motors consist of the portal, large terminase (TerL) and small terminase (TerS) subunits, each of which assembles into a homomeric ring (19,20). Genome packaging by terminases can be broadly summarized as a five step process (21) (Figure 1). (Step 1) First, the motor recognizes the concatemeric viral genome, primarily through TerS binding (22–27). (Step 2) Next, TerL cleaves the DNA at a specific site and binds to the portal complex. (Step 3) TerL uses ATP hydrolysis to translocate DNA (28–30) through the portal ring into the capsid (31). (Step 4) Upon completing the translocation of at least one genome-length of DNA, TerL switches its enzymatic activity from translocation to cleavage (32). This cleavage occurs either after encapsidating exactly one genome length (termed ‘unit-length packaging’) or after the capsid is completely filled with DNA, resulting in slightly more than one-genome length being packaged (termed ‘headful packaging’). (Step 5) Finally the terminase subunits are released from the capsid for maturation of another virus, while portal binds to the tail proteins to complete a mature, infectious virion (13). Although the sequence of these events has been well studied, the structural mechanism for each step is largely unknown. In particular, how the motor holds DNA during either translocation or cleavage remains obscure.

**Figure 1:**
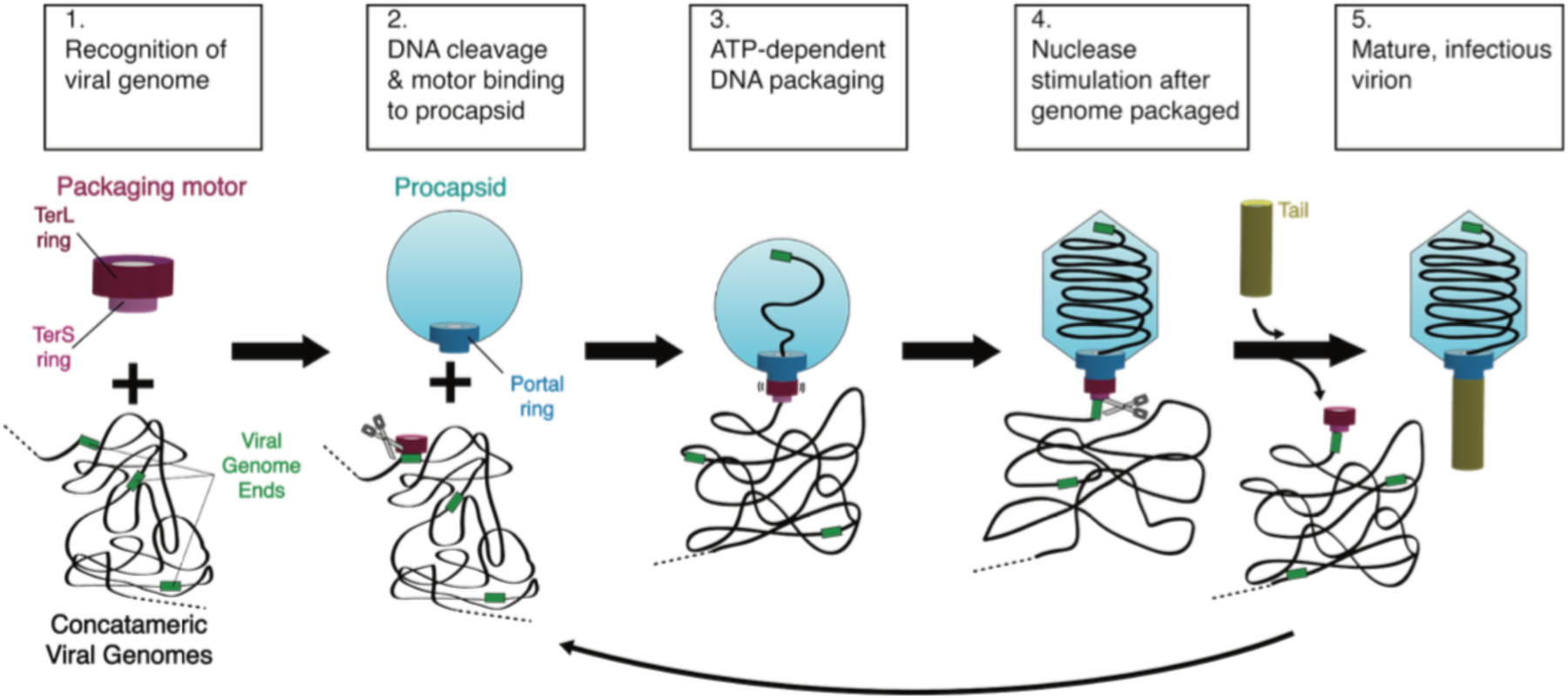
Schematic of a generic genome packaging reaction catalyzed by a terminase enzyme.

The TerL protein is the catalytic engine of the packaging apparatus, harboring the two enzymatic activities of the motor: the ATPase that drives DNA translocation, and the endonuclease that cleaves genome concatemers at both initiation and termination of packaging (33). The terminase motor is capable of generating high force (stall force up to ~60 pN) and high speeds (up to ~2000 bp/s) (2,34). However, several distinct structural mechanisms have been proposed for both the force generation reaction and nucleolysis (7,18,35–41). Even more mysterious is the mechanism of nuclease activity regulation; current models for nuclease regulation include auto-inhibition by of TerL (38,42), catalytic regulation via competition between fast DNA translocation and slow nucleolysis (37,43), and inhibition mediated by TerS (37,39,42,44). Careful dissection of TerL structure and mechanism is necessary to discern between competing models for TerL activity and regulation.

TerL contains two domains: a C-terminal nuclease domain of a RNaseH fold (7,18,35,38,39,41) and an N-terminal ATPase domain of the ASCE (<u>a</u>dditional <u>s</u>trand, <u>c</u>onserved <u>g</u>lutamate) superfamily (45). The TerL protein forms a pentameric ring when bound to the procapsid (35). The C-terminal tail of TerL is thought to interact with the portal and/or the procapsid (33,46–52), although an alternate arrangement in which the N-terminal ATPase domain contacts the portal has also been proposed (35,53). Our group recently proposed a model for the TerL ring in which the ATPase domains form a tight ring with intersubunit contacts contributing to ATP hydrolysis and a small inner pore for binding DNA (52). In this model, the nuclease domains are positioned on the periphery of the TerL ring so that they can use their C-terminal tails to interact with the portal ring. Because the ATPase domains line inner pore of the ring, this model suggests that the ATPase domains are the primary point of contact for DNA during DNA translocation. However, this model did not explain how DNA cleavage occurs. How does the nuclease domain contribute to DNA binding? Does the interaction surface change during the cleavage reaction?

Here we report enzymatic and structural characterization of TerL DNA binding and nuclease function. We use TerL from the thermophilic phage P74-26 (TerL^P74-26^) (54) due to its high expression, solubility, and stability. Employing a thermophilic terminase affords us a unique opportunity to separately evaluate DNA binding and nucleolytic cleavage, as the latter function only occurs at elevated temperatures. We show that both tight DNA binding and cleavage are nucleotide dependent. Our analysis of cleavage kinetics reveals that dual strand cleavage is fast, suggesting that multiple nuclease domains collaborate to simultaneously cut the double helix. We also report the structure of the P74-26 nuclease domain, which we use to map the contributions of individual residues to both DNA binding and cleavage. Our data indicate that the ATPase domain is the primary determinant of DNA binding and that the nuclease domain is dispensable for DNA binding. We integrate our results to propose a mechanism for how TerL switches between DNA translocase and nuclease modes.

## Materials and Methods

### Protein expression and purification

Both the isolated ATPase domain (1–256) and full-length P74-26 TerL mutants were expressed and purified as previously described (52). The nuclease domain (residues 256–485) was subcloned from our previously described pet24a full-length construct and was overexpressed identically to the above constructs (52). Cells were lysed in a cell disruptor and pelleted. The lysate was applied to a 10-mL His-Trap column (GE Healthcare) preequilibrated in buffer A (500 mM NaCl, 20mM Imidazole, 50 mM Tris pH 8.5, 5 mM βME, 10% glycerol). The column was washed with buffers A, followed by buffer A’ (150 mM NaCl, 20 mM Imidazole, 50 mM Tris pH 7.5, 5 mM βME, 10% glycerol). Protein was eluted with buffer B (150 mM NaCl, 250 mM Imidazole, 50 mM Tris pH 8.5, 5 mM βME, 10% glycerol). Eluate was dialyzed into buffer QA (125 mM NaCl, 25 mM Tris pH 7.5, 2 mM DTT, 10% glycerol) and the tag was cleaved with prescission protease overnight. Dialysate was loaded onto a 10-mL Q column (GE Healthcare) preequilibrated with buffer QA. The column was then washed with buffer QA. Protein was eluted by applying a 0–100% gradient of buffer QA to buffer QB (1 M NaCl, 25 mM Tris pH 7.5, 2 mM DTT, 10% glycerol). Eluate was injected onto an S200 HR26/60 (GE Healthcare) column preequilibrated with gel filtration buffer (125 mM NaCl, 25 mM Tris pH 7.5, 4 mM DTT). Eluted protein was concentrated to ~20 mg/mL and flash frozen in liquid nitrogen.

### Crystallization, structure determination, and refinement

Native crystals formed in hanging drops containing 20 mg/mL TerL Nuclease domain mixed 2:1 with buffer containing 0.23 M sodium phosphate monobasic/potassium phosphate dibasic pH 6.2 and 2.5 M sodium chloride, and 4 mM dTMP. Crystals were plunged into cryoprotectant containing 0.28 M sodium phosphate monobasic/potassium phosphate dibasic pH 6.2, 4 M sodium chloride, and 2.5 mM dTMP before being flash frozen in liquid nitrogen. Data were collected at the Advanced Light Source at SIBYLS beamline 12.3.1 at wavelength 1.000 Å. Heavy atom derivative crystals were obtained by incubating native crystals with 3 mM potassium hexachloroplatinate 24 hours prior to flash freezing. Cryoprotectant for heavy atom derivative crystals contained 3.4 mM potassium hexachloroplatinate. Derivative crystal data were collected at the Advanced Photon Source GM/CA CAT beamline 23ID-B at wavelength 0.855 Å in inverse beam mode. All diffraction data were processed with HKL3000 (55). Platinum bound to Met 265 allowed SAD phasing (56) of the 2.7 Å derivative crystal dataset using the PHENIX autosol pipeline (57). Native dataset anisotropic diffraction data were corrected with the UCLA Diffraction Anisotropy Server (58) and phases were extended to 2.6 Å resolution. Model building and structure refinement were performed with COOT (59) and PHENIX (60). The structure was then deposited in the RCSB (PDB code 5TGE).

### DNA binding and Nuclease digestion

ADP-Beryllium Fluoride (ADP-BeF_3_) was formed by incubating 50 mM Tris pH 8.5, 150 mM potassium chloride, 1 mM DTT, 1 mM ADP, 10 mM sodium fluoride, 4 mM beryllium chloride, and 10 mM magnesium chloride for two hours prior to usage. TerL and ADP-BeF_3_ were mixed and incubated for 5 minutes prior to addition of 150 ng of plasmid pET28a. Upon DNA addition, samples were incubated at room temperature (DNA binding) or 60° C (nuclease digestion) for 30 minutes unless otherwise indicated. During kinetics experiments, addition of cold 25 mM EDTA (final) and rapid cooling in an ice bath quenched cleavage. Unless otherwise noted, all DNA cleavage samples were quenched with 1.5% SDS (final) to prevent TerL’s DNA binding activity from perturbing DNA migration through the gel. Standard 1.5% agarose Tris-Acetate EDTA pH 8.0 gels were used, with the exception of isolated ATPase domain DNA-binding assay. The pH of the running buffer and gel were raised to pH 8.5 to account for the ATPase domain PI of 8.1. Gels were imaged on an LAS 3000. Gel densitometry was performed using ImageJ (61). Kinetic modeling was performed using the Curve Fitter module in Berkeley Madonna (62). Berkeley Madonna does not report fitted errors. However, the fits to each model were extremely robust as the initial values of the fitted parameters could be set up to 10-fold higher or lower than their actual values, and ultimately the same results were produced. (If initial values were set 100-fold higher or lower, the fitting routine could not converge on a reasonable solution.)

### Isolation of P23-45 Phage Genomic DNA

An initial stock of phage P23-45 was kindly provided by the Severinov laboratory. A fresh culture of *Thermus thermophilus* HB8 was grown to an OD_600_ of ~1.0 in growth medium (0.8% (w/v) Tryptone, 0.4% (w/v) Yeast Extract, 0.3% (w/v) NaCl, 1 mM MgCl_2_, and 0.5 mM CaCl_2_) (63). 150 µL of fresh culture was combined with 100 µL P23-45 phage stock at a concentration of 10^6^ Plaque Forming Units per mL (PFU/mL) and incubated at 65°C for 10 minutes. This mixture was then inoculated into 20 mL of fresh growth medium and incubated for 4–6 hours at 65°C. The culture was spun at 4,000 × g for 20 minutes to remove cell debris. Supernatant (>10^9^ PFU/mL) was then treated with DNase I (final concentration, 2 Units/mL) and incubated at 30°C for 30 minutes. Genomic DNA was extracted from P23-45 phage stocks using the Phage DNA Isolation Kit (Norgen Biotek Corp) according to the manufacturer’s protocol.

## Results

### TerL^P74-26^ displays robust nuclease activity

We previously established that TerL^P74-26^ binds DNA at room temperature in the presence of ADP•BeF_3_ (52). Because P74-26 phage is a thermophile, we hypothesized that the relatively low temperature (~20 °C) of our previous DNA binding experiments ‘masked’ the underlying endonuclease activity. Indeed, raising the temperature activates TerL^P74-26^’s nuclease activity, with robust cleavage at ~40 °C that accelerates to maximal cleavage at 60 °C (Figure 2A). TerL^P74-26^ also requires to be locked in an‘ATP-bound’ state by a non-hydrolyzable ATP analog to efficiently cleave DNA (Figure 2B). Indeed, we only observe DNA cleavage with TerLP74-26 pre-loaded with ADP•BeF_3_ and no cleavage in the apo or ADP-loaded states. We observe no significant cleavage by TerL^P74-26^ in the presence of ATP, because robust TerL ATPase activity (52) rapidly depletes all available ATP. Therefore, TerL^P74-26^ needs to be locked in an ‘ATP-bound’ state for productive cleavage, indicating a strong linkage between the ATPase and nuclease activities.

**Figure 2:**
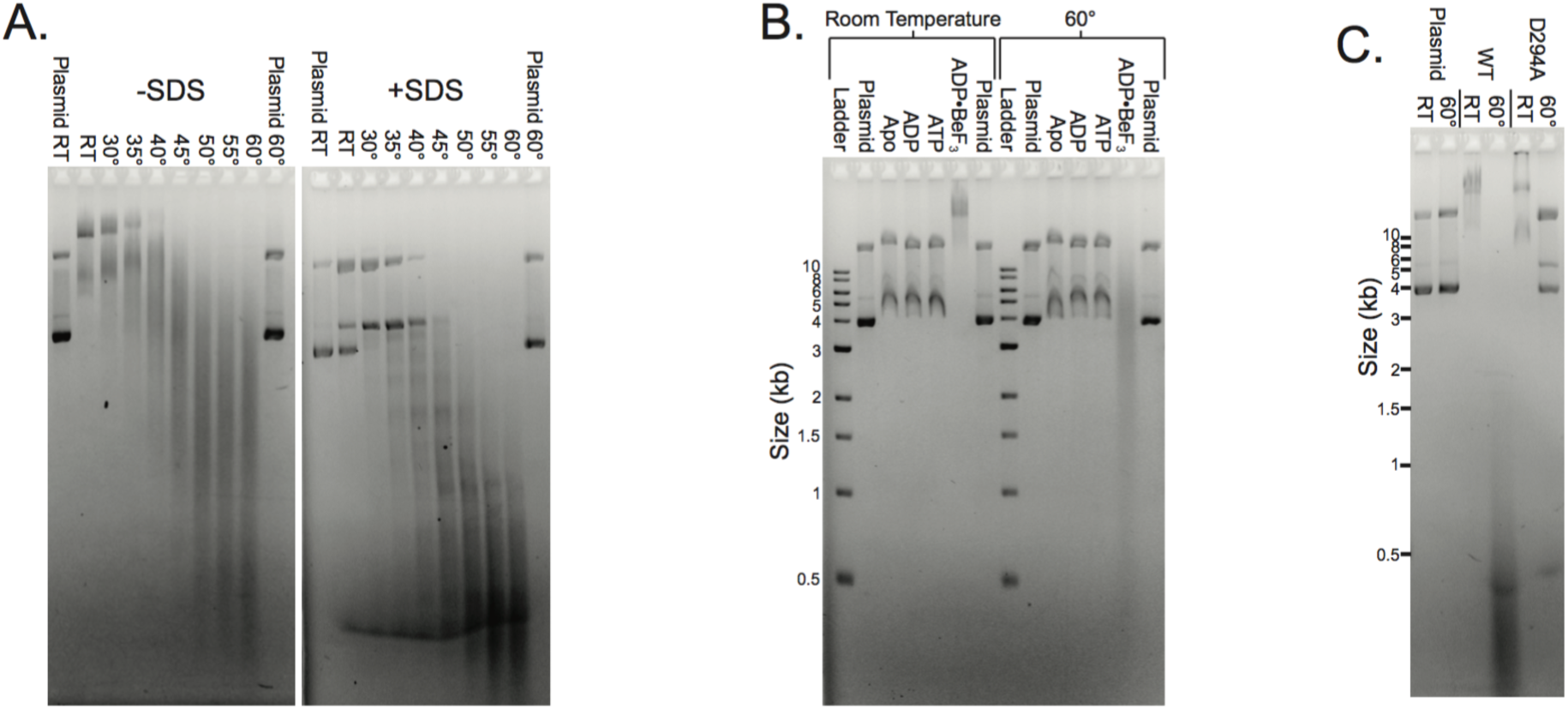
Characterization of TerL^P74-26^ DNA-binding and nucleolytic activity. (A) Elevated temperature activates TerL^P74-26^ nucleolysis. Plasmid DNA migrates slowly when mixed with 15 μM TerL^P74-26^ (left panel, no SDS) at low temperatures, indicating TerL^P74-26^ primarily binds DNA at these temperatures. DNA cleavage occurs at higher temperatures indicated by the low molecular weight smearing. Addition of SDS (right panel) reveals only minimal cleavage at room temperature (RT) with increased intensity of the nicked and linear plasmid bands. At temperatures ≥40 °C, we observe robust cleavage that increases as the temperature is raised. (B) At room temperature, DNA weakly binds to TerL^P74-26^ in the apo state or when incubated with ADP or ATP. Locking TerL into an “ATP-bound” state with the nonhydrolyzable analog ADP•BeF_3_ results in tight DNA binding. At 60 °C TerL cleaves DNA, but only in the presence of ADP•BeF_3_. Buffer control samples containing ADP•BeF_3_ do not exhibit perturbed plasmid migration (final lane at each temperature). (C) Mutation of D294, the conserved nuclease active site residue necessary for metal binding metal binding residue, to alanine results in a severe loss of nucleolysis (10 μM protein) while not affecting TerL^P74-26^’s affinity for DNA.

To exclude the possibility of a co-purified contaminant being responsible for the observed nucleolysis, we mutated an absolutely conserved metal-coordinating active site residue (D294) to alanine. TerL^P74-26^ is expected to have a two-metal coordinated active site as is found in related nucleases (7,37–39,42,64,65). D294 is the best candidate for mutagensis because this residue coordinates one of the metal ions in the active site (7,35,37–39). The D294A variant displays an almost complete loss of cleavage activity, while retaining tight DNA binding affinity (Figure 2C). These results illustrate that the nucleolytic activity is due to TerL^P74-26^ and not a contaminant.

Therefore, isolated TerL^P74-26^ retains the three critical activities necessary for terminase function: ATPase activity, DNA-binding activity (52), and DNA cleavage.

To our knowledge, TerL^P74-26^ is the only known large terminase that can bind and cleave DNA as a full-length protein. The isolated ATPase domain of TerL^T4^ is competent to bind DNA, whereas the full-length protein shows no significant affinity for DNA (66). Other full-length TerL proteins exhibit significant *in vitro* nuclease activity (7,37,39,41,67), although DNA-binding activity is undetectable. Therefore both DNA binding and cleavage can be separately dissected with TerL^P74-26.^

We next investigated specificity of the DNA binding and cleavage activities of TerL^P74-26^. As we have shown previously, TerL^P74-26^ binds DNA with no sequence specificity, as all bands in a 1kb-ladder are shifted (52). When incubated at 60° C, TerL^P74-26^ degrades all bands in the 1kb-ladder, as well as negatively supercoiled plasmid, linearized plasmid, and other linear fragments (Supplemental Figure 2). Thus, TerL^P74-26^ both binds and cleaves DNA with no discernible sequence specificity. TerL proteins from other phage cleave DNA with no sequence specificity, indicating that TerL^P74-26^ is similar to most other TerL proteins in this respect.

To investigate the linkage between DNA binding and cleavage, we measured the TerL^P74-26^ concentration dependence for both activities. DNA binding and cleavage reactions were performed at different temperatures, but otherwise with identical reaction conditions and an identical plasmid substrate. Supercoiled plasmid allows for facile measurement of single- or dual-strand cleavage during nuclease activity assays. Strikingly, we observe that DNA binding and DNA cleavage exhibit nearly identical dependence on TerL concentration (Figure 3A-B). At TerL^P74-26^ concentrations ≤2 µM, no DNA binding or cleavage is observed within the 30 minutes of the reaction. However, at >2 µM, TerL^P74-26^ exhibits significant binding at room temperature, with higher TerL^P74-26^ concentrations resulting in slower DNA migration. Likewise, at 60 °C TerL^P74-26^ concentrations higher than 2 µM result in substantial fragmentation of the plasmid over the 30-minute time course. Increasing incubation time to 16 hours at 60 °C results in substantial, but incomplete DNA cleavage at 2 µM TerL^P74-26^, with minor cleavage at 1.5 µM (Supplemental Figure 3). The coincident TerL dependencies of DNA binding and cleavage activities indicate that these two functions are tightly linked. Moreover, the steepness of the activity transition suggests that a cooperative process drives both DNA binding and cleavage. The affinity of TerLP74-26 for DNA (Kd ~3 µM) is similar to that measured for the Lambda phage TerL protein (Kd ~3–4 µM) (68), indicating that TerL^P74-26^ is consistent with known terminase enzyme function.

**Figure 3:**
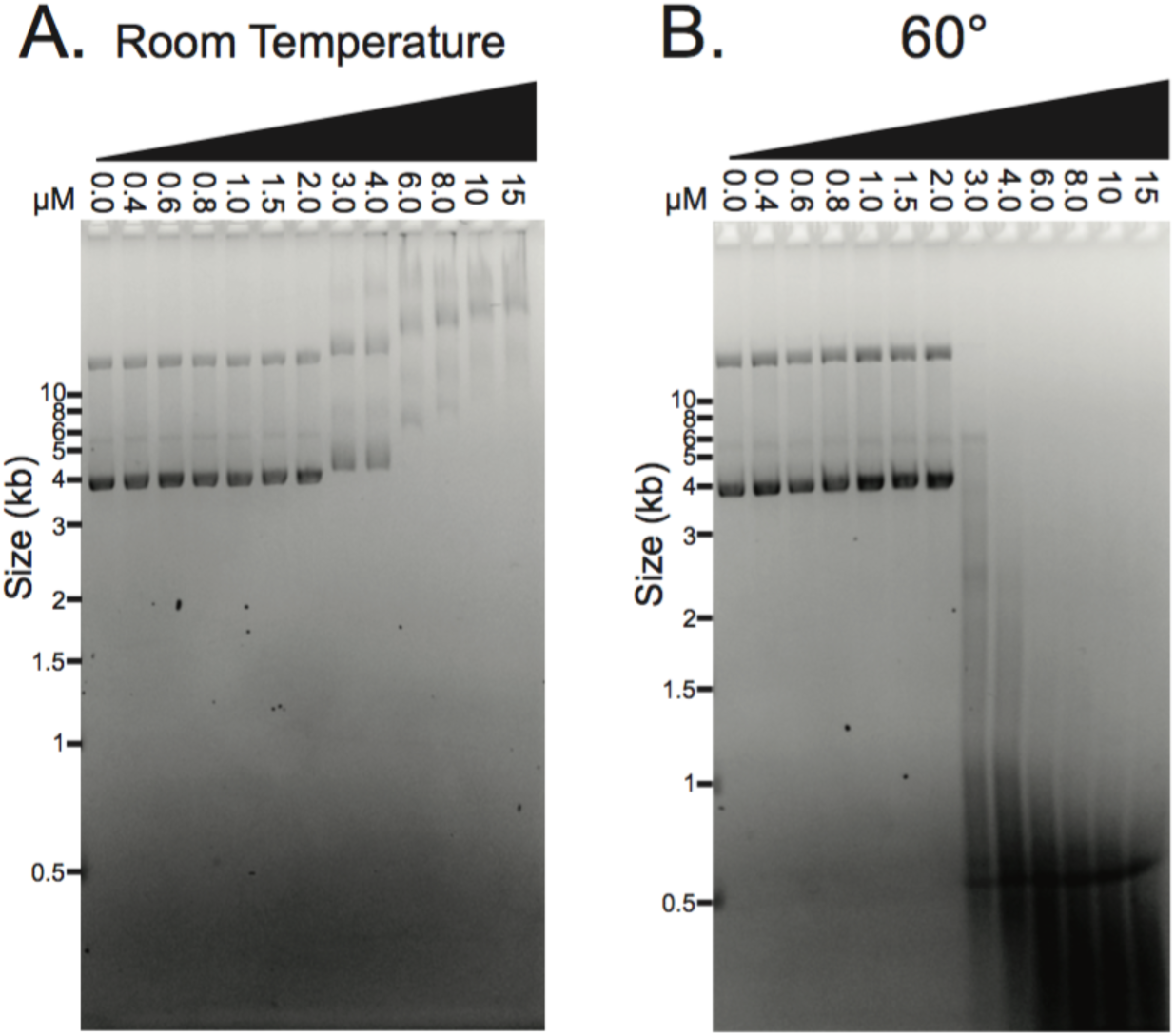
DNA binding and cleavage exhibit identical protein concentration dependence. (A) At room temperature, full length TerL^P74-26^ binds DNA at concentrations greater than 2 μM (no SDS added to the samples). DNA migrates slower, indicating greater binding, as the concentration of TerL^P74-26^ is raised. (B) At 60 °C, TerL^P74-26^ cleaves DNA proportional to the degree of DNA binding observed at room temperature. Tighter binding (slower migration) at room temperature corresponds with more complete DNA digestion at 60 °C. SDS was used to remove any TerL bound to DNA prior to loading samples.

### Kinetic Analysis of TerL^P74-26^ cleavage

To further investigate the mechanism of TerL-mediated nucleolysis, we followed the kinetics of plasmid cleavage. We chose a TerL^P74-26^ concentration of 5 μM due to the potent activity observed during the 30-minute reaction. By measuring the intensities of the supercoiled, nicked, and linear plasmid bands over a 10-minute time course, we can distinguish between single-strand versus dual-strand cleavage. To accurately quantify each band, we added SDS to our gel-loading buffer to prevent TerL^P74-26^’s DNA binding activity from perturbing DNA migration. We observe a rapid loss of supercoiled DNA ^(t1/2~20 sec), whereas the nicked and linearized plasmid bands increase and then^ decrease in intensity (Figure 4A). The increase in nicked plasmid (peak ~30 seconds) roughly matches the decrease in supercoiled plasmid. Nicked plasmid is then degraded until it is undetectable at ~480 seconds. Linearized DNA increases until ~60 seconds, and then is degraded to form a smear, indicating more substantial degradation.

**Figure 4:**
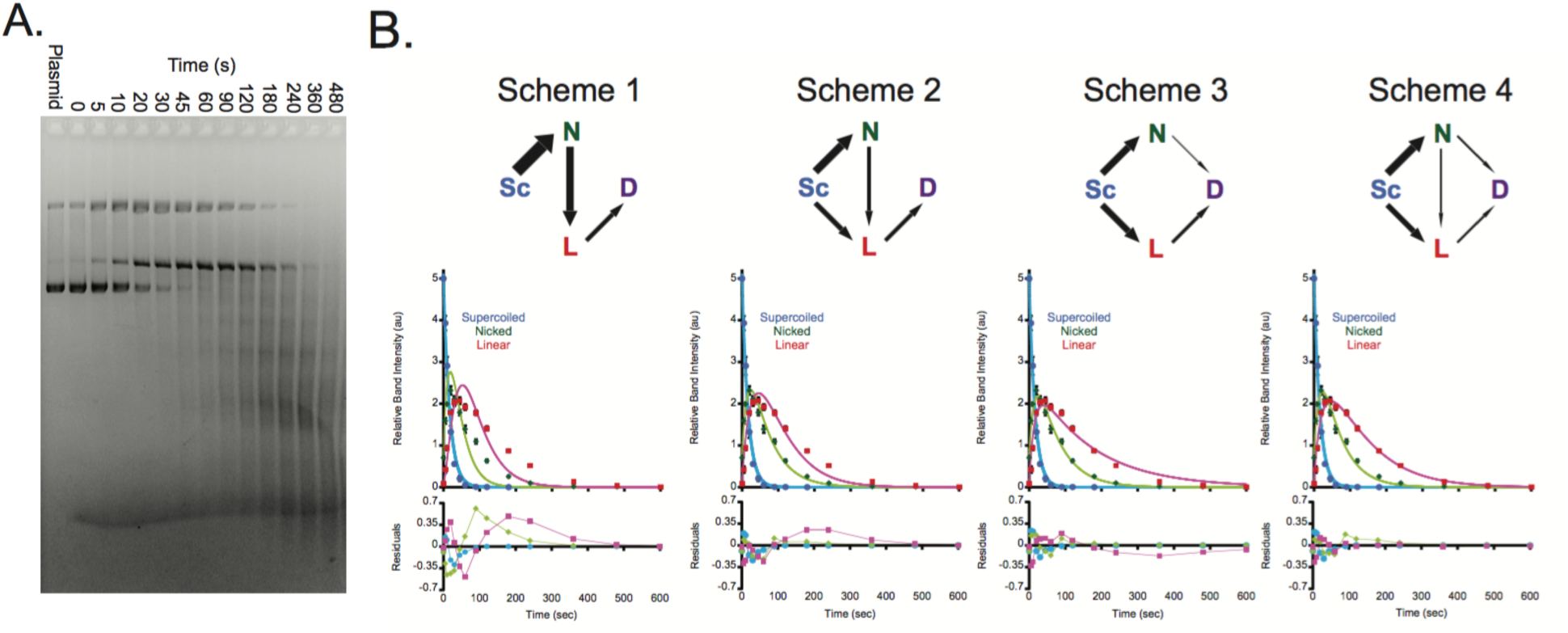
Kinetics of TerL^P74-26^ nuclease activity. (A) A representative gel with 5 μM TerL^P74-26^ incubated with plasmid at 60 °C for different durations. Densitometry of replicates measures band intensity for the plasmid’s supercoiled, nicked, and linear bands to calculate cleavage rates and create kinetic models. (SDS present in loading buffer.) (B) Kinetic models fit to the kinetic data (four replicates). Four schemes each represent different models. Arrows indicate reaction pathways between supercoiled (Sc), nicked (N), linearized (L), and digested (D) species. Arrow thickness is scaled to the value of the calculated rate (Supplemental Table 1). The fit of each scheme is indicated below the reaction pathway. The residuals of the fit for each species are plotted at the bottom to show the quality of the fit for each scheme. Higher quality fitting requires simultaneous cleavage of both DNA strands.

Because we observe rapid plasmid cleavage, we sought to determine whether a more physiologically relevant substrate would exhibit substantially different kinetics. A digestion time course with the genome of related phage P23-45 (92% identical between nucleotide sequence of the P23-45 and P74-26 genomes; 99.8% identity between amino acid sequence of P23-45 and P74-26 TerL proteins (63)) showed a similar rate of digestion (Supplemental Figure 4). Therefore, we hypothesize that regulation of TerL nuclease activity originates from factors other than DNA sequence.

We used kinetic modeling to test different possible schemes for the nucleolysis reaction. Kinetic schemes were tested in Berkeley Madonna (62) by fitting rate constant values to experimental data. The simplest potential mechanism to describe the TerL^P74-26^ nuclease reaction is that TerL first cleaves one strand of DNA to produce a nicked plasmid, followed by cleavage of the opposing strand to generate a linear product, which can then be fully degraded (Figure 4B, Scheme A). This straightforward nucleolytic mechanism (with three fitted parameters) has been assumed for other TerL cleavage reactions (7,41). Surprisingly, this reaction scheme does not fit the data well, as the rise and fall of the nicked and linearized plasmid bands are fundamentally distinct from that predicted by this model (note the clear non-random distribution of the residuals). Therefore, we explored other potential mechanisms for the nucleolysis reaction. Interestingly, only reaction schemes involving simultaneous dual-strand cleavage reasonably fit the data (Figure 4, Schemes B-D). In scheme B, TerL can either nick or directly linearize the plasmid; nicked plasmid then becomes linearized before full degradation can proceed. Scheme B, which has four separate fitted parameters, improves the fit relative to scheme A, but still exhibits significant deviation from the data. In scheme C, TerL can either nick or directly linearize the plasmid, and each of these intermediates is degraded to small fragments. In scheme D, TerL can either nick or directly linearize the plasmid. The nicked intermediate can be either converted to a linearized DNA or directly degraded into small fragments. Schemes C and D match the data best, as indicated by examining the residuals of the fit (Figure 4B). Because Scheme D has one more fitted parameter than Scheme C (5 vs. 4), it is difficult to confidently distinguish between the two schemes.

Regardless, our kinetic modeling approach indicates that TerL is capable of cleaving both strands simultaneously or near simultaneously. The rate of nicking (Supplemental Table 1) is nearly identical to that of linearization (Scheme 4; k_nicking_ = 0.037 s^−1^, k_linearization_ = 0.026 s^−1^). The comparable rates for nicking and linearization suggest that plasmid linearization is not the result of two completely separate nicking reactions, but rather simultaneous (or near simultaneous) dual-strand cleavage. This result has important impacts on how the nuclease functions in the TerL protein (see Discussion).

### Structure of TerL^P74-26^ nuclease domain

To gain insight into the structural mechanism of nucleolysis by TerL^P74-26^, we solved the structure of the TerL^P74-26^ nuclease domain (hereafter, TerL^P74-26^-ND) to 2.6 Å resolution (Figure 5A). We obtained experimental phases from single-wavelength anomalous diffraction with a platinum derivative (see Materials & Methods) (Table 1). The overall fold of TerL^P74-26^ -ND is similar to those of other terminase nuclease domains (7,35,38–41), with an average C_α_ RMSD of 2.0 Å (for individual C_α_ RMSDs, see Supplemental Table 2).

**Table 1.** Crystallographic statistics of the TerL^P74-26^-ND structure

| Data Collection |  |
| --- | --- |
| Space Group | P 43 21 2 |
| Wavelength | 1.00 |
| Resolution Range | 47.51-2.60 |
| Unit Cell | 71.37 71.37 127.32 |
| Unit Cell | 90 90 90 |
| Total Reflections | 299519 |
| Unique Reflections | 10480 (862) |
| Multiplicity | 28 (28.8) |
| Completeness % | 98 (98) |
| Mean I/sigma I | 23.3 (6.8) |
| Wilson B factor | 31.9 |
| R-merge | 0.089 (0.638) |
| R-meas | 0.090 (0.650) |
| R-pim | 0.017 (0.121) |
| CC1/2 | 0.999 (0.999) |
| CC* | 1.000 (0.999) |

| Refinement |  |
| --- | --- |
| R-work % | 21.8 |
| R-free % | 24.7 |
| RMS Bonds | 0.003 |
| RMS Angles | 0.52 |
| Ramachandran Favored % | 94 |
| Ramachandran Outliers % | 0 |
| Rotamer Outliers % | 0.62 |
| Clashscore | 4.05 |
| Average B | 38 |

Several high-resolution structures of the large terminase nuclease domain for highly related phage, G20C, are solved in the accompanying article from Xu et al. The protein sequences for TerL^G20C^-ND and TerL^P74-26^-ND are nearly identical, and differ only at residue 315 (G20C A315 vs P74-26 V315). TerL^G20C^-ND crystallized in three crystal forms that are distinct from TerL^P74-26^-ND. Overall, the structures of TerL^G20C^-ND and TerL^P74-26^-ND complement one another to provide key insight into TerL nuclease structure and function.

**Figure 5:**
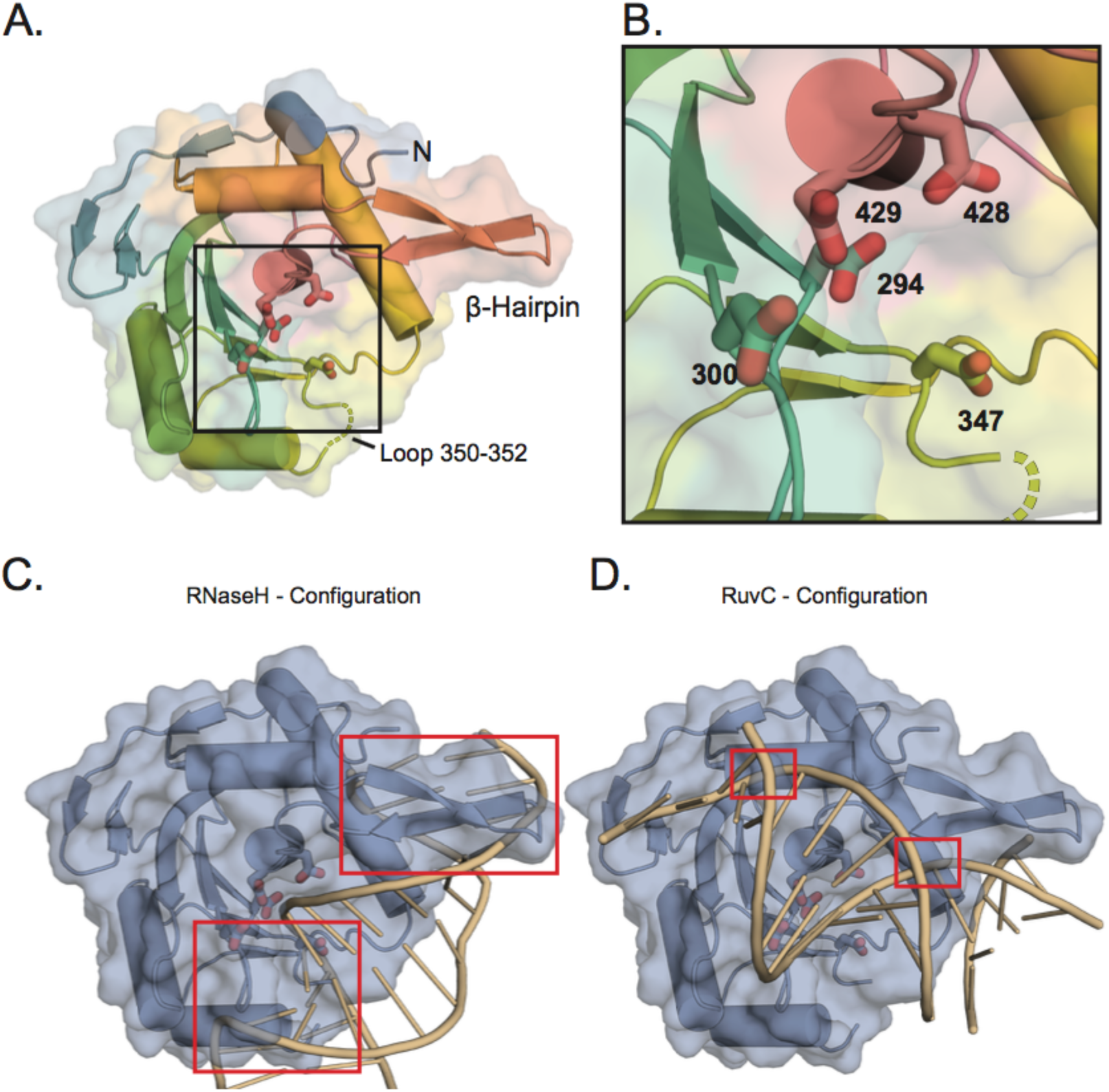
The TerL^P74-26^ nuclease domain structure. (A) Overall features of the TerL^P74-26^ nuclease domain structure. The electron density is missing for both a flexible loop at position 350–352 (Gly-Val-Gly; dotted lines) and for C-terminal residues 450–485. Potential metal-coordinating active site residues (294, 300, 347, 428, and 429) are represented with sticks. A β-hairpin that is unique to the terminase family extends away from the nuclease domain. (B) Zoomed view of the nuclease active site shows the acidic residues for metal coordination. (C) The RNase H configuration of DNA bound the TerL^P74-26^-ND was created by aligning an RNase H structure bound to a RNA/DNA hybrid duplex (70) to the TerL^P74-26^-ND using default parameters in Chimera (76). The RNA:DNA duplex clashes (red boxes) with TerL^P74-26^-ND. There are severe clashes in the regions around metal-coordinating residue 347 and the β-hairpin. (D) The RuvC configuration of DNA bound to TerL^P74-26^-ND was generated using the structure of RuvC resolvase bound to a Holliday junction (71). Clashing is minimal, occurring at G424 and the side chains of R421 and R425. The flexibility of these residues suggests they may change conformation to accommodate DNA binding.

The active site of TerL^P74-26^ -ND contains several metal-coordinating residues that are conserved across the terminase family. Because no divalent cations were added during purification and crystallization, we do not observe any metal coordination in the active site. As expected from structures of other TerL nuclease domains (7,35,38–41), we observe D294 in the heart of the active site accompanying several other metal-coordinating residues (D294, D300, D347, D428, D429) in TerLP74-26 (Figure 5A-B). The positions of acidic residues in TerL^P74-26^ most closely resembles the arrangement observed in phage T4/RB49 (Supplemental Figure 5), which binds metal with residues equivalent to TerL^P74-26^ D294, D347, and D429 (35,37). The accompanying article by Xu et al. discusses metal coordination in detail for TerL^G20C^-ND.

Beyond the active site residues, there is remarkably little sequence conservation within the nuclease domain across the TerL family. The relative lack of conservation may reflect the fact that different viruses use varied strategies for cleaving DNA (21,69). Because of this lack of sequence conservation, identifying how DNA accesses the nuclease active site has been particularly challenging.

To address how DNA is positioned in the TerL nuclease active site, we compared the TerL nuclease domain to other, more distantly related nuclease structures for which there are structures of substrates bound. In particular, the structure of human RNaseH bound to an RNA:DNA hybrid (70) and a more recent structure of *T. thermophilus* RuvC resolvase bound to a Holliday junction (71) provide two different possibilities for the DNA orientation in the nuclease active site. By superposing these two structures with that of TerL^P74-26^-ND, we can model potential DNA interaction modes. Superposition of the RNaseH:RNA-DNA structure positions the DNA running from a flexible loop at residues 350–352, across the active site towards the β-hairpin and N-terminus of the nuclease domain (Figure 5C). However, as has been noted previously (38,39,41), this positioning clashes with a β-hairpin that is present in all TerL proteins but is absent in other known members of the RNaseH superfamily of nucleases (38). Therefore, in the RNaseH configuration, the β-hairpin must exhibit flexibility in order to accommodate DNA. Because of this substantial clash, the β-hairpin has been proposed to play an auto-regulatory role in controlling nuclease activity (38). In contrast, superposition of RuvC suggests an orthogonal DNA orientation, with the DNA running across the active site in a directions roughly parallel to the β-hairpin (Figure 5D). Importantly, the β-hairpin does not produce a significant clash with modeled DNA but would instead provide a surface for cradling the DNA as it crosses the active site. Thus, the two RNaseH and RuvC models predict different roles for the β-hairpin: the RuvC model predicts that the β-hairpin assists in DNA cleavage while the RNaseH model predicts that the β-hairpin inhibits cleavage.

To further investigate the role of the nuclease domain in terminase function, we used the TerL^P74-26^ nuclease structure to identify residues that may be important for DNA binding and nucleolysis. We selected conserved or semi-conserved basic residues that are predicted to contact DNA, based on previous predictions of DNA binding surfaces (7,18,35,38,39,41) and our comparisons with RNaseH and RuvC. Combined with variants in the ATPase domain that we previously generated to study ATP hydrolysis and DNA binding (52), our panel includes 23 point mutations across both domains (Supplemental Table 3). We also used our isolated TerL^P74-26^ ATPase domain (TerL^P74-26^-AD) and TerL^P74-26^-ND constructs to examine the overall role of each domain in TerL function. By separately measuring DNA-binding and nuclease cleavage for each variant, we provide critical insight into how DNA is bound and cleaved during viral genome packaging.

### The ATPase domain is the primary DNA binding region

To assess how DNA binds to TerL^P74-26^ we first focused on the ATPase domain, as we previously observed a complete loss of DNA binding from the R101E mutation (52). R101 is in a patch of basic residues along one surface of the ATPase domain that we predict forms the DNA binding surface within the pore of the assembled TerL ring (52). Here we extend this analysis to other residues across the surface of the ATPase domain, including other residues in the ‘basic patch’. Three of the mutations in the basic patch (R102A, R104E, and R128A) also display a complete loss of DNA binding, while the final basic patch variant (R121E) does not significantly affect binding (Figure 6A and Supplemental Figure 6A-B). To verify that these DNA-binding effects are due to specific disruption of the DNA binding interface, we tested variants with mutations predicted to be outside of the pore of the TerL ring (R58A and R170A). Neither of these mutations severely affects DNA binding affinity (Figure 6A). These results support our previous conclusion that this basic patch is critical for gripping DNA.

**Figure 6:**
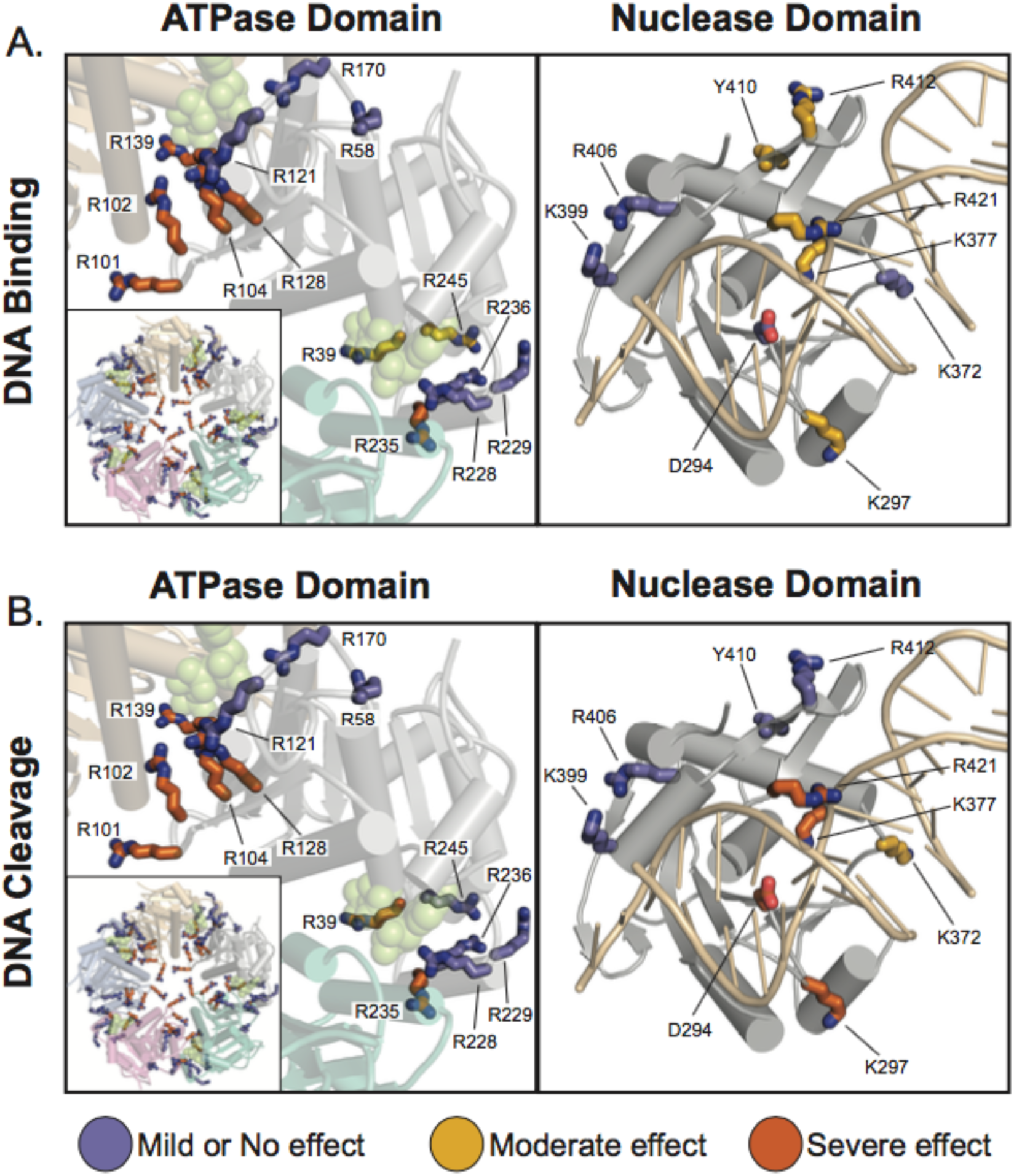
Mapping of residues important for DNA binding and cleavage. (A) Mutation of ATPase domain (left panel) basic patch residues severely inhibits DNA binding whereas nuclease domain variants do not severely impact binding (right panel). DNA binding for each variant was assessed based on how far the most intense bands migrated into the gel. The ATPase domain is shown in the context of our previous model of the ATPase ring (52) to illustrate how mutational effects match the ring topology (green spheres indicate ADP•BeF_3_ in the ATPase active site). The nuclease structure is shown with DNA in the RuvC configuration. Mutations that severely inhibit binding (orange) tend to be in the pore of the ATPase ring. Nuclease domain variants fail to significantly inhibit DNA binding. (B) Mapping of residues important for DNA cleavage. Similar to the binding assessment, variants were ranked by their ability to efficiently cleave DNA. Residues in the ATPase domain that are important for DNA binding are likewise critical for DNA cleavage (left panel). DNA binding is therefore a prerequisite for effective cleavage. Residues that are predicted to interact with DNA in the RuvC configuration R297, K377, and R421) are critical for nuclease activity, suggesting that DNA binds similarly to RuvC. Nuclease metal coordinating variant D294A serves as a control.

Because TerL^P74-26^ needs to be locked into an ATP-bound state to tightly grip DNA (Figure 2B), we next investigated how mutations in or near the ATPase active site affect DNA binding. R39 is a conserved residue in the P-loop of the active site and directly contacts the γ-phosphate group of ATP (52). R139 is the *trans*-acting arginine finger that is critical for ATP hydrolysis (52). We also tested several residues (R228, R229, R235, R236, R245) in the Lid subdomain, a region that caps the active site and changes conformation upon ATP hydrolysis and release (Figure 6A) (52). The R228A, R229A, and R236A variants have no apparent effect on DNA binding, while the R39A and R245A variants exhibit a moderate decrease in DNA binding. Only the R139A and R235A variants display severe defects in DNA binding. Because R139 and R235 are important for both ATP hydrolysis and interfacial contacts between ATPase subunits (52), we hypothesize that the DNA binding defects observed with these variants is due to the severe loss of both ATP binding and ring assembly.

We next focused on the role of the nuclease domain in DNA binding. We mutated residues in the active site (D294A and K377A), the β-hairpin (Y410A, R412A, and R421E), and other regions predicted to bind DNA (K297A, K372A, K399A, and R406A). Interestingly, none of the mutations in the nuclease domain severely impact DNA binding (Figure 6A). Variants K297A and K377A display a moderate loss of affinity for DNA. Similarly, the mutations in the β-hairpin display modest defects. Neither R412 nor R421 are conserved in the β-hairpin, but structures of terminase nuclease domains often exhibit basic residues in similar locations (35,38–40), suggesting that basic residues may play some role in function. A third β-hairpin mutation (Y410A), designed to disrupt the β-hairpin structure, only moderately affected DNA binding. Overall these results indicate that the nuclease domain is not a primary determinant for high affinity DNA binding.

Because our panel of point mutants highlights the importance of the ATPase domain in binding DNA, we next investigated whether isolated domains bind DNA. TerL^P74-26^-AD binds DNA at similar concentrations as full-length TerL^P74-26^ (Figure 7A and Supplemental Figure 7A). Interestingly, TerL^P74-26^-AD binds DNA independent of nucleotide, indicating that the nuclease domain is important for the ATP-dependent regulation of DNA binding. In contrast, TerL^P74-26^-ND does not detectably bind DNA, even at concentrations >30-fold higher than the Kd for full-length TerLP74-26 binding (Figure 7B). Therefore, the ATPase domain is necessary and sufficient for TerL to bind DNA, an event that is a prerequisite for nuclease activity.

**Figure 7:**
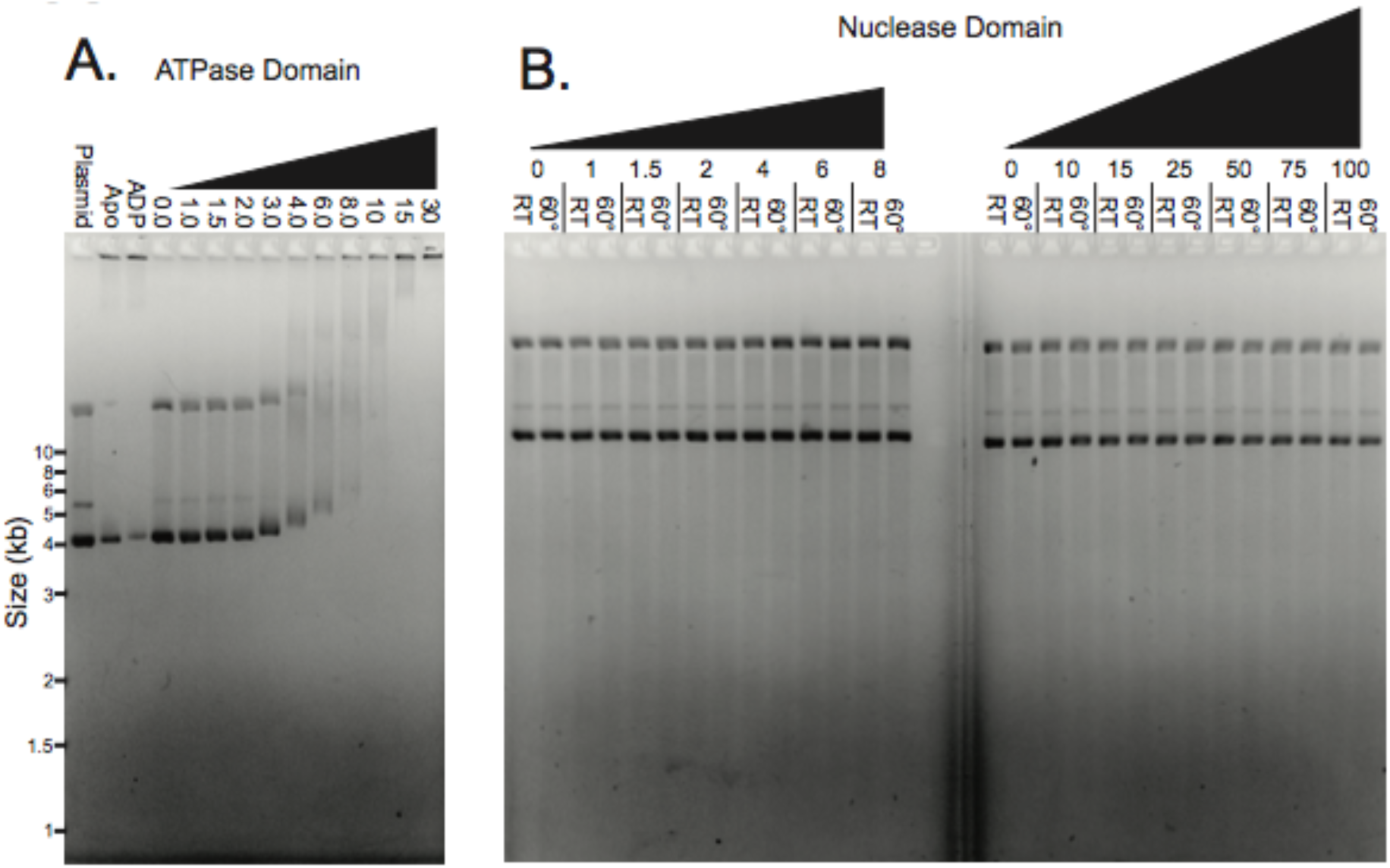
The ATPase domain is necessary and sufficient for DNA binding. (A) The isolated TerL^P74-26^-AD binds DNA with the same affinity as full length TerL^P74-26^ independent of ADP•BeF_3_. 15 μM TerLP74-26-AD slows migration of a significant portion of plasmid DNA in the apo form or the presence of ADP. In the presence of ADP•BeF_3_, DNA migration slows with TerL^P74-26^-AD concentrations above 2 μM with decreasing migration proportional to the rise in protein concentration. Coomassie staining confirms the presence of TerL^P74-26^-AD co-migrating with DNA (Supplemental Figure 6). (B) The isolated TerL^P74-26^-ND neither binds nor cleaves DNA, even at concentrations 30-fold higher than where we observe binding and cleavage for full-length TerL^P74-26^.

### Identifying the requirements for nucleolysis

We next examined our panel of variants to determine the role of individual residues on the nucleolysis reaction. We find a strong correlation between DNA binding and nucleolysis across all ATPase mutants. Mutations that abrogate DNA binding likewise inhibit nuclease activity, while mutations in the ATPase domain that do not disrupt DNA binding have no effect on nuclease activity. Specifically, mutations in the ATPase domain’s basic patch (R101E, R102A, R104E, and R128A) or the active site (R39A, R139A, R235A) show a severe loss of nuclease function (Figure 6B and Supplemental Figure 6C). These results support our finding that the isolated TerL^P74-26^-ND fails to bind and cleave DNA. Therefore binding and nucleolysis hinge on the ATPase domain’s ability to bind DNA.

Interestingly, a subset of nuclease domain mutants disrupts nucleolysis without severely impairing DNA binding. We observe a severe loss of nuclease activity in the K297A, K377A, and R421E variants, and a moderate decrease in activity in the K372A variant. K297A, K377A, and R421E are the only variants in TerL^P74-26^ where DNA binding remains relatively unperturbed yet nucleolysis is severely impacted. As mentioned previously, the RuvC and RNaseH binding modes predict very different behavior for several of the variants, particularly those with mutation in the β-hairpin. The RuvC binding mode predicts a favorable role for the β-hairpin in nucleolysis, while the RNaseH mode predicts that the β-hairpin plays an auto-inhibitory role. Overall, cleavage defects in these variants are consistent with the DNA contacts predicted by the model of DNA bound to TerL^P74-26^-ND based on the RuvC structure (Figure 5D). In particular, R421 is on the face of the β-hairpin predicted to interact favorably with DNA in the RuvC binding mode; this residue is necessary for DNA cleavage activity (Figure 6B). In contrast, Y410 and R412 are on the opposite face of the β-hairpin and are dispensable for nucleolysis. Regardless of whether the RuvC or RNaseH binding modes are correct for TerL nuclease engagement, our results demonstrate that cleavage depends on two factors, 1) the ability of TerL to bind DNA as dictated primarily by the ATPase domain, and, 2) a specific set of nuclease domain residues predicted to position DNA for nucleolysis.

## DISCUSSION

Proper terminase function in viral genome packaging requires precise spatiotemporal coordination and regulation of ATP hydrolysis, DNA binding, and nucleolytic cleavage. Each of these functions must be individually examined in order to piece together a packaging mechanism. We previously built a low-resolution structural model of the pentameric TerL ring that accurately predicted the position of the arginine finger (R139 in TerL^P74-26^), as well as a key DNA-binding residue (R101 in TerL^P74-26^) (52). We observed ATP-dependent conformational changes in the ATPase domain, suggesting that the Lid subdomain generates the force for DNA translocation through a lever-like motion. In this study we provide insights into terminase function that can then be applied to improve the existing models of genome packaging and further our understanding of one of nature’s most powerful bio-motors.

### The TerL ATPase domain tightly grips DNA

The TerL ATPase domain is indispensable for DNA binding. This conclusion is based on two major observations. First, we observe strong DNA binding with both full-length TerL^P74-26^ and the isolated ATPase domain. In contrast, the isolated nuclease domain of TerL^P74-26^ does not bind or cleave DNA. Thus, the ATPase domain is both necessary and sufficient for DNA binding. Second, mutation of basic patch residues (R101, R102, R104, or R128) in full-length TerL^P74-26^ abrogates both DNA binding and nucleolytic cleavage. Conversely, none of the mutations located in the nuclease domain severely impact DNA binding, despite the fact that several residues are critical for nucleolysis. These results suggest that the bulk of TerL affinity for DNA derives from the ATPase domain, as predicted by our previous model (52).

Several models for terminase:DNA binding predict a larger role of the nuclease domain in gripping DNA during translocation (18,35). Because we can separately measure DNA binding and cleavage with TerL^P74-26^, we are able to directly test this prediction. Surprisingly we find that residues within the nuclease domain only make a small contribution to DNA affinity (Figure 6A) and that the entire domain is dispensable for tight binding (Figure 7A and Supplemental Figure 7). Moreover, two semi-conserved residues (K399 and R406) in a region predicted to bind DNA (35) show no role in binding or cleaving DNA (Figure 6A). Therefore, we favor a model in which the ATPase domain is the primary DNA grip during both translocation and cleavage modes, and that the nuclease domain only engages DNA during genome cleavage (Figure 8). Although unlikely, it is possible that free TerL^P74-26^ is locked into ‘cleavage mode’ and uses the nuclease domain for gripping DNA when in ‘translocation mode’. However, we do not favor this model because we observe no measurable affinity between the nuclease domain and DNA (Figure 7B), similar to results seen with T4-TerL (66). Moreover, the isolated ATPase domain, which is unlikely to be locked into ‘DNA cleavage mode’, displays tight DNA binding, as shown here (Figure 7A) and elsewhere for TerL^T4^ (66).

**Figure 8:**
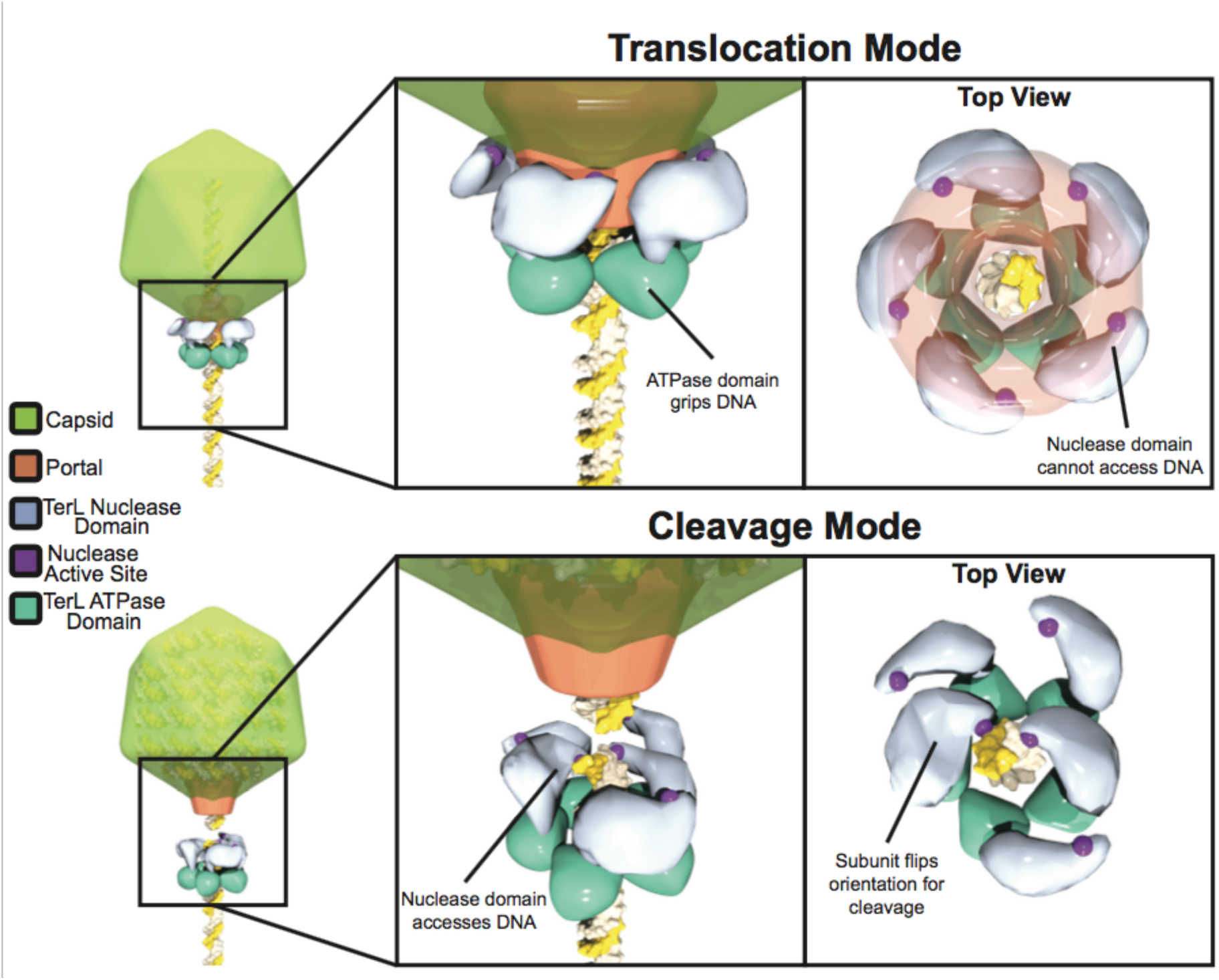
Proposed model for nuclease regulation. During ‘translocation mode’ the nuclease domain active site is sequestered from DNA by interactions of the TerL with portal and capsid, preventing premature cleavage. The ATPase domain serves as the sole surface for gripping DNA during packaging. Upon completion of packaging TerL enters ‘cleavage mode’. TerL dissociates from the portal and capsid, releasing the inhibition of the nuclease domains. The ATPase domains remains tightly bound to DNA. The nuclease domains rearrange to simultaneously cleave each of the antiparallel DNA strands. Although depicted as a blunt cut, cleavage could also leave overhangs depending on how both nuclease domains engage DNA.

By separating the primary DNA gripping region from the nuclease active site, terminases have evolved an efficient means for regulating nuclease activity. First, the nuclease domain’s low intrinsic DNA-binding affinity appears to be critical for proper nuclease regulation. Although the nuclease active site must bind DNA with at least weak affinity in order to cleave, this affinity must be carefully balanced; TerL would catalyze spurious cleavage if the affinity were too strong, but would cleave inefficiently if the affinity were too weak. There appears to be a spectrum of intrinsic DNA binding affinities for TerL nuclease domains. Isolated T4- (37,66), Sf6- (18,40), CMV- (7), and HSV-TerL (41) nuclease domains cleave DNA, implying a modest affinity. On the other hand, isolated SPP1- (38), P22- (39), and P74-26-TerL nuclease domains fail to cleave DNA. Secondly, the flexible nature between the ATPase and nuclease domains allows the allosteric regulation of the nuclease. The ATPase domain places the nuclease domain in high local concentration with DNA to overcome the nuclease domain’s intrinsically weak affinity for DNA. Moreover, by altering the position of the nuclease active site relative to DNA, terminase enzymes can easily regulate DNA cleavage (see below).

### Kinetic modeling reveals details of TerL nucleolytic activity

Kinetic analysis of the TerL^P74-26^ nuclease activity reveals a mechanism of TerL cleavage consistent across the terminase family. TerL^P74-26^ rapidly cleaves supercoiled plasmid, with concomitant increases in nicked and linearized plasmid, followed by complete digestion. Qualitatively similar results were observed for T4 phage and CMV TerL-catalyzed nucleolysis (7,42). Similarly, increasing concentrations of HSV-1 TerL nuclease domain results in an initial increase in nicked and linearized plasmid, followed by near complete digestion at high nuclease concentrations (41). These results were interpreted as single strand nicking that eventually results in linearization and complete degradation (7,41). In contrast, our kinetic modeling of the nucleolysis reaction reveals significant simultaneous dual strand cleavage by TerL^P74-26^. This result places large constraints on the arrangement of the TerL nuclease domains during nucleolysis. Because the qualitative cleavage data is consistent across the family, we propose that these constraints are universal to all terminases.

From a mechanistic perspective, simultaneous dual strand cleavage requires flexibility of the nuclease domain relative to the ATPase domain. Endonucleases require two active sites arranged in an anti-parallel fashion for simultaneous dual strand cleavage (72,73). As previously suggested (37,42), the TerL nuclease domains would assume conformations roughly 180° relative to one another for their active sites to align with each of the antiparallel DNA strands. Therefore, there is significant flexibility between the ATPase and nuclease domains of TerL to allow this rearrangement. Indeed, there is much previous data to support this assertion. Limited proteolysis of the TerL proteins from P74-26 (Supplemental Figure 8), T4 (33) and P22 (39) indicates that the linker connecting the ATPase Lid subdomain to the nuclease domain is highly flexible. Additionally, crystal structures of TerL proteins from T4 and Sf6 show very different orientations of the nuclease domain relative to the ATPase domain (18,35). Therefore, we propose that the TerL ATPase domain ring tightly grips DNA while the TerL nuclease domain is flexibly tethered to adopt the necessary orientation for DNA cleavage.

How does dual strand cleavage occur? We envision two possibilities for cleavage of both strands: 1) a monomer of TerL cleaves both strands in rapid succession, or 2) two subunits within a TerL oligomer cleave each strand simultaneously or near simultaneously. In the first mechanism, after cleaving the Watson strand, the nuclease domain of the TerL monomer must rapidly reorient by ~180° to cleave the Crick strand. The second cleavage reaction would have to be faster than the time scale of our experiments, because our kinetic modeling shows that two sequential strand cleavage events do not occur within the same time scale as each other. In the second mechanism, two separate nuclease domains within a TerL oligomer can adopt orthogonal orientations to efficiently cleave both strands. Although we cannot decisively rule out dual strand cleavage by a monomer, we favor cleavage by a TerL oligomer for two reasons. First, the steep dependence on TerL concentration for both DNA binding and cleavage implies assembly of a TerL oligomer on DNA (Figure 3). Second, TerL requires ATP for both DNA binding and cleavage (Figure 2B), which implies that the interfacial contacts afforded by ATP binding (52) promote oligomerization on DNA. Regardless, both mechanisms require a large degree of flexibility between the tightly bound ATPase domain and the nuclease domain.

### Regulation of TerL nuclease activity

During translocation TerL nuclease activity must be inhibited to prevent premature cleavage. Two non-mutually exclusive possibilities could explain how TerL nuclease activity is regulated: 1) ‘kinetic competition model’, the rate of TerL ATP hydrolysis and DNA translocation significantly outpaces the rate of nucleolysis until translocation slows upon maximal packaging, allowing cleavage to occur, and 2) ‘steric block model’, procapsid components and/or TerS regulate the accessibility of the nuclease active site for DNA. We discuss these two possibilities below.

In the kinetic competition model the relative rates of ATP hydrolysis and nucleolysis self-regulate TerL cleavage (37,43). If DNA translocation is much faster than the rate of nucleolysis, then the nuclease active site cannot stably engage DNA long enough for cleavage. Packaging would progress until the rate of DNA translocation sufficiently slows near the end of packaging. As the rates of translocation and nucleolysis become similar, the nuclease domain has enough time to engage a segment of DNA for successful cleavage. In the kinetic competition model, the rates of ATP hydrolysis and nucleolysis must be precisely balanced to prevent premature genome cleavage. However, two lines of evidence suggest that kinetic competition is not regulation mechanism. First, motor stalling events regularly occur during packaging in phages T4 (2) and Lambda (34,74) for periods of time up to ~ 5 seconds with no reported cleavage of DNA. Second, long-term motor stalls can been artificially induced in phage T4 with no significant cleavage occurring over the time scale of hours (49,75).

In the steric block model, inhibition is achieved by restricting the accessibility of the nuclease active site for DNA. Multiple lines of evidence indicate that the TerL C-terminal tail binds to the portal (33,46–52). We propose that this interaction locks the nuclease domain in an orientation that prevents the nuclease active site from accessing DNA, thereby preventing premature genome cleavage. We further hypothesize that dissociation of TerL from portal upon completion of packaging releases the nuclease domain from this restricted conformation, allowing the flexibly tethered nuclease domains to reorient and doubly cut DNA. TerS can inhibit TerL-nuclease activity (37,39,42,44); therefore, similar contacts with TerS could sterically block TerL nuclease activity.

How does DNA engage the nuclease active site? We use both the structure of RNaseH bound to an RNA/DNA hybrid (70) or RuvC bound to a Holliday junction (71) to model how DNA accesses the active site. DNA binds in orthogonal orientations in these structures, leading to predictions for behavior of some of the variants tested here. The phenotypes of the D294A, K297A, and K377A (important for cleavage) and K399A and R406A (no role in cleavage) variants match both the RuvC and RNaseH orientations and do not effectively discriminate between the two models. We favor a model of DNA binding similar to RuvC. TerL is more similar to RuvC than RNaseH in terms of structure and the surface of TerL is a better steric fit to the RuvC DNA orientation rather than that of RNaseH. As noted here and in other studies (38,39), the β-hairpin would clash with DNA in the RNaseH-like configuration (Figure 5C). These results have raised the question of whether the β-hairpin adjusts its conformation to allow for productive access to the active site or if DNA is bound in a different orientation to the RNaseH model. In contrast, the model based on RuvC does not result in any clash between the DNA and the β-hairpin (Figure 5D). Instead, the β-hairpin is positioned such that it can make favorable interactions along the DNA backbone. Importantly, our mutagenesis results are most consistent with the orientation of DNA predicted from the RuvC model. A β-hairpin residue predicted to directly form a salt bridge with the DNA backbone of the scissile strand (R421) is critical for nuclease activity, but two residues on the opposite face of the hairpin (Y410 and R412) are dispensable. In additional to R421, three other residues in our panel of mutations are also predicted to interact with the scissile strand in the RuvC-like model (K297, K377, and R412); all three are critical for nuclease activity (Figure 6B). Furthermore, K372 is predicted to interact with the non-scissile strand, and the K372A variant shows a modest defect in nucleolytic activity. Future studies will map the interactions with DNA in greater detail.

Terminases are conserved across many different families of dsDNA viruses, including human pathogens of Herpesviridae (11). There are several FDA-approved drugs on the market that target the terminase motor, and it is thought that these drugs’ mode-of-action is through inhibition of the terminase’s nucleolytic activity (3–9). Our studies reveal important aspects of the TerL nuclease mechanism and regulation, and provide a blueprint for future mode-of-action studies of small molecule inhibitors of the terminase.

## Acknowledgements

The authors would like to thank the Severinov laboratory (Rutgers Univ.) for providing a sample of phage P23-45. We thank the Schiffer, Royer, and Ryder labs for use of instrumentation and for helpful discussions. We thank members of the Antson Lab (University of York) for sharing data before publication and for thoughtful discussions. We thank beamline scientists at SIBYLS 12.3.1 (Lawrence Berkeley National Laboratory) and APS 23-ID-B (Argonne National Laboratory, award number GUP-39936) for technical support with x-ray diffraction data collection. B.A.K. is a Pew Scholar in the Biomedical Sciences, supported by the Pew Charitable Trusts.

## Supplemental Figures

**Supplemental Figure 2:**
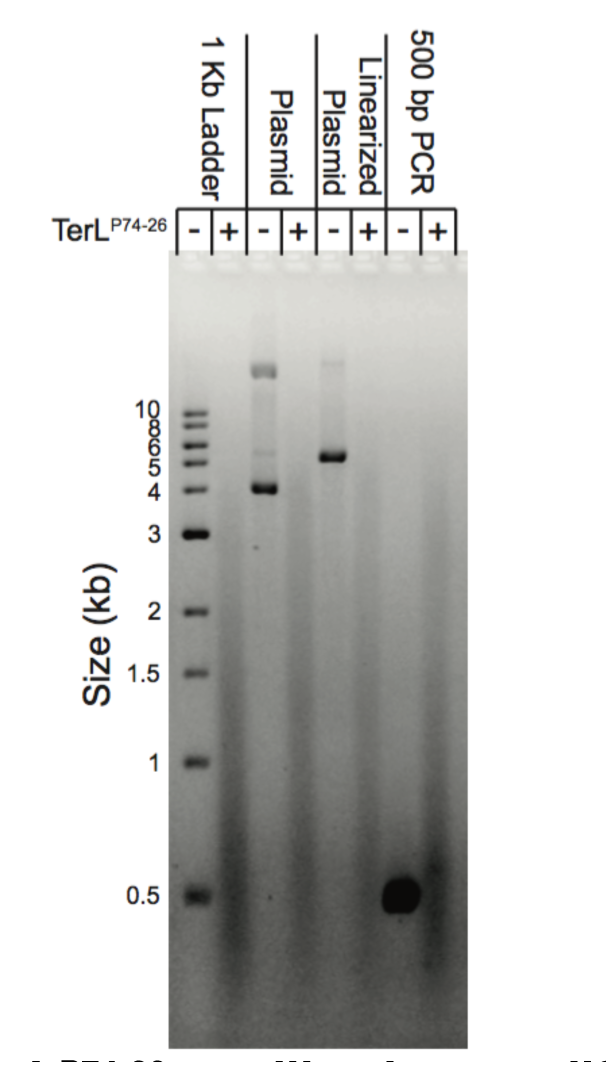
TerL^P74-26^ readily cleaves different DNA substrates. Untreated substrates are compared to substrates incubated with 15 μM TerL^P74-26^. TerL^P74-26^ cleaves a 1 kb ladder, plasmid, linearized plasmid, and a 500 bp PCR product. No SDS was used in the loading dye resulting in diffuse smearing due to digested products remaining bound to TerL^P74-26^.

**Supplemental Figure 3:**
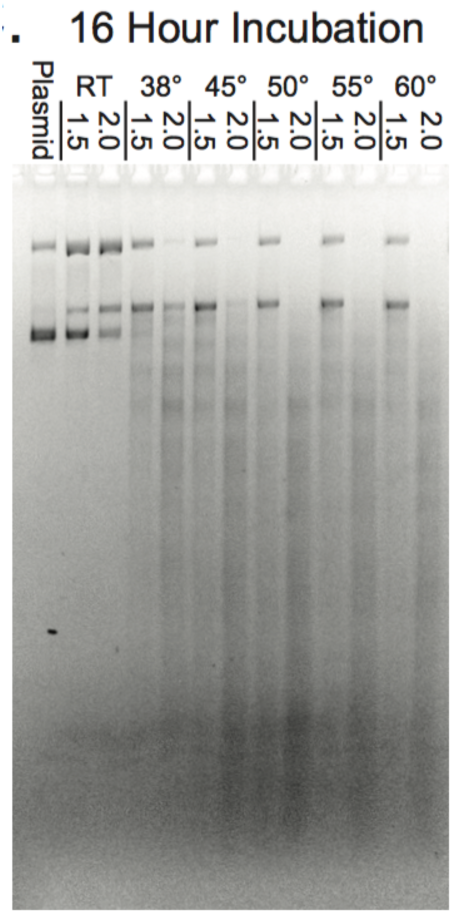
At TerL^P74-26^ concentrations where we do not observe binding in 30 minutes (1.5 and 2 μM), long incubation (16 hours) results in limited DNA cleavage across various temperatures. SDS was not present in loading dye for this gel.

**Supplemental Figure 4:**
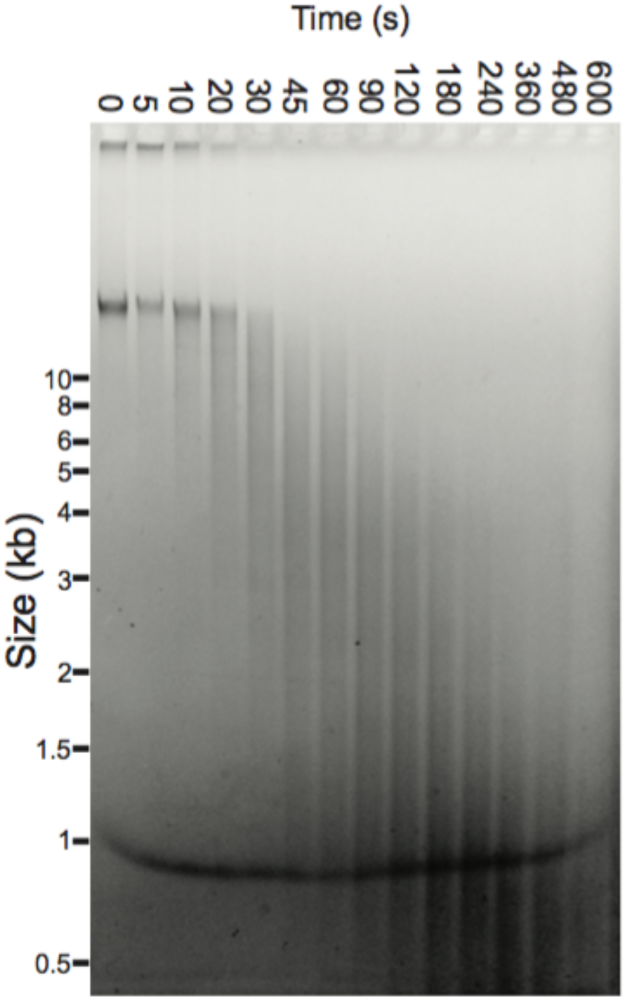
Purified phage P23-45 genome (100 ng) digested using the same time course used for kinetic analysis of plasmid cleavage. The genome is rapidly cleaved indicating the genome sequence does not regulate cleavage.

**Supplemental Figure 5:**
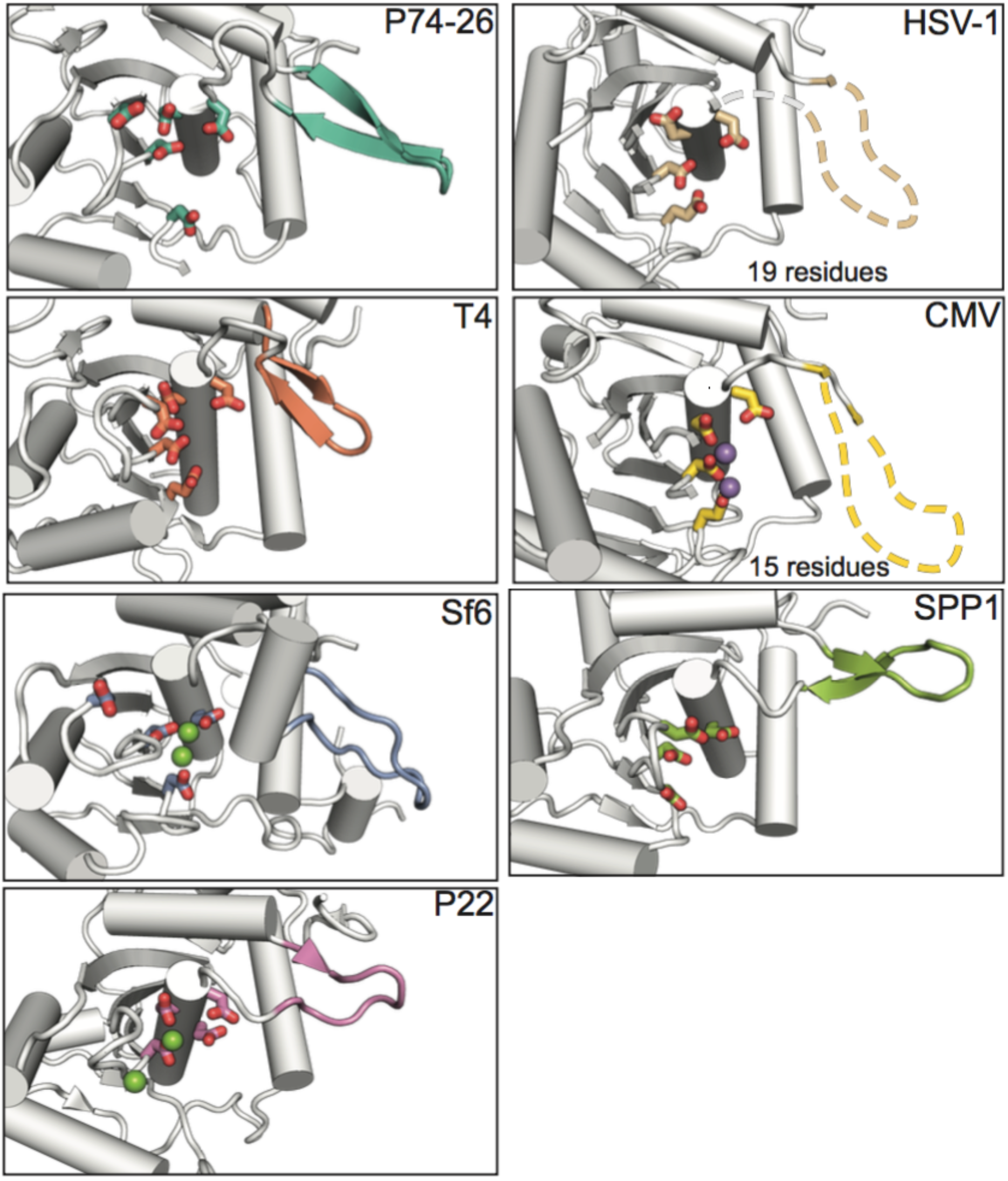
Structural comparison of the nuclease domain active site and β-hairpin across terminase family: TerL^P74-26^, T4 (35), Sf6 (40), P22 (39), SPP1 (38), HSV-1 (41), and CMV (7). Sticks represent potential or confirmed metal coordinating residues. Bound Mg^2+^ (green spheres) or Mn^2+^ (purple spheres) are present in Sf6, P22, and CMV structures. The β-hairpin (colored main chain) is variable in size across the terminase family. The two structures of herpesviruses TerL-ND (from HSV-1 and CMV) have large disordered β-hairpin regions (dotted lines). The remaining structures have variably sized β-hairpins. The P74-26 active site most closely resembles that of configuration of acidic residues observed in the T4 structure.

**Supplemental Figure 6:**
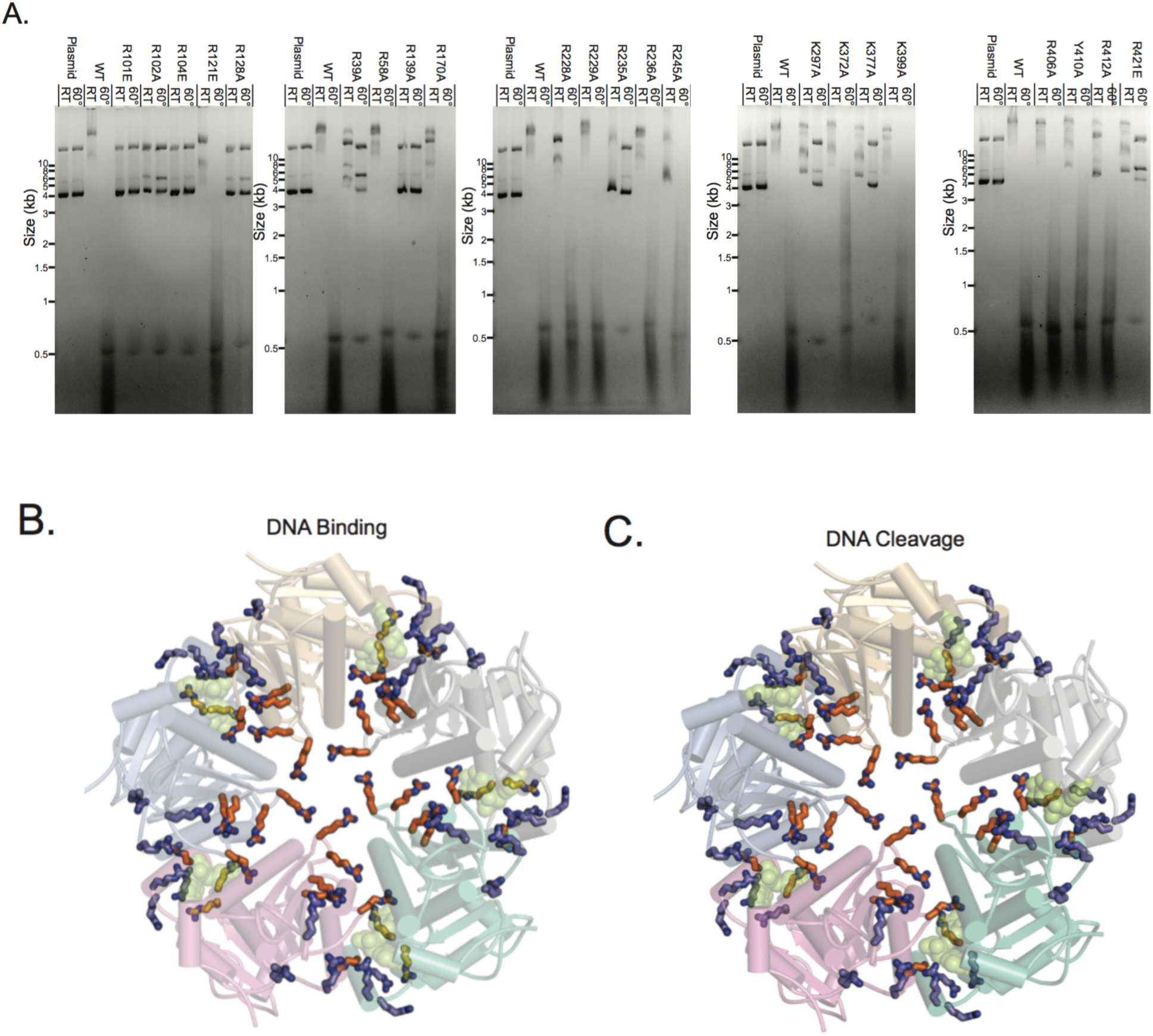
(A) 10 μM of each variant was assessed for ability to both bind and cleave DNA. These mobility shift gels were then used to rank the variants. SDS was not included in the loading dye for room temperature samples. (B & C) The insets of the Figure 6 (left panels) are enlarged to enhance the view of the effects of the variants on (B) DNA binding, and (C) DNA Cleavage in the context of the ATPase ring model (52).

**Supplemental Figure 7:**
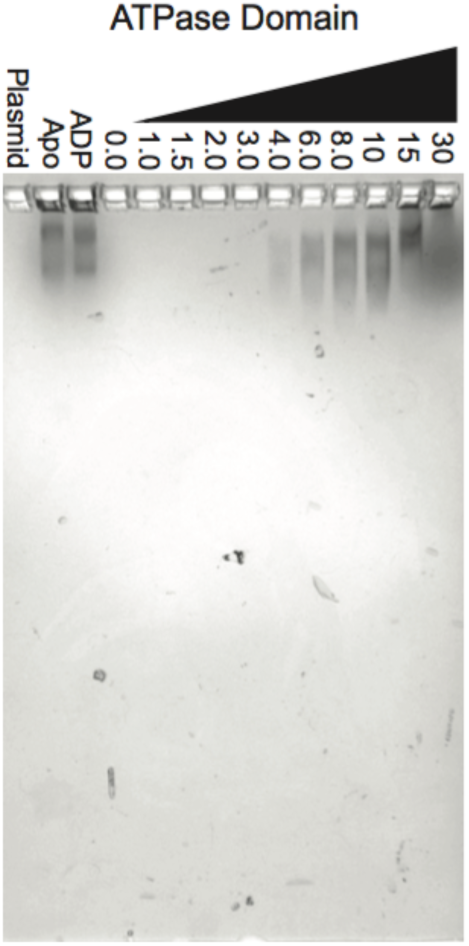
Coomassie staining of the agarose gel from Figure 6A shows TerL^P74-26^-AD binds DNA. The theoretical PI of TerL^P74-26^-AD is 8.1, preventing significant migration of naked protein into the gel, resulting in staining at TerL^P74-26^-AD bound DNA or at the border of the gel lane wells. The pH of the gel (8.5) was modified slightly to account for the TerL^P74-26^-AD PI without perturbing binding to DNA (Materials and Methods).

**Supplemental Figure 8:**
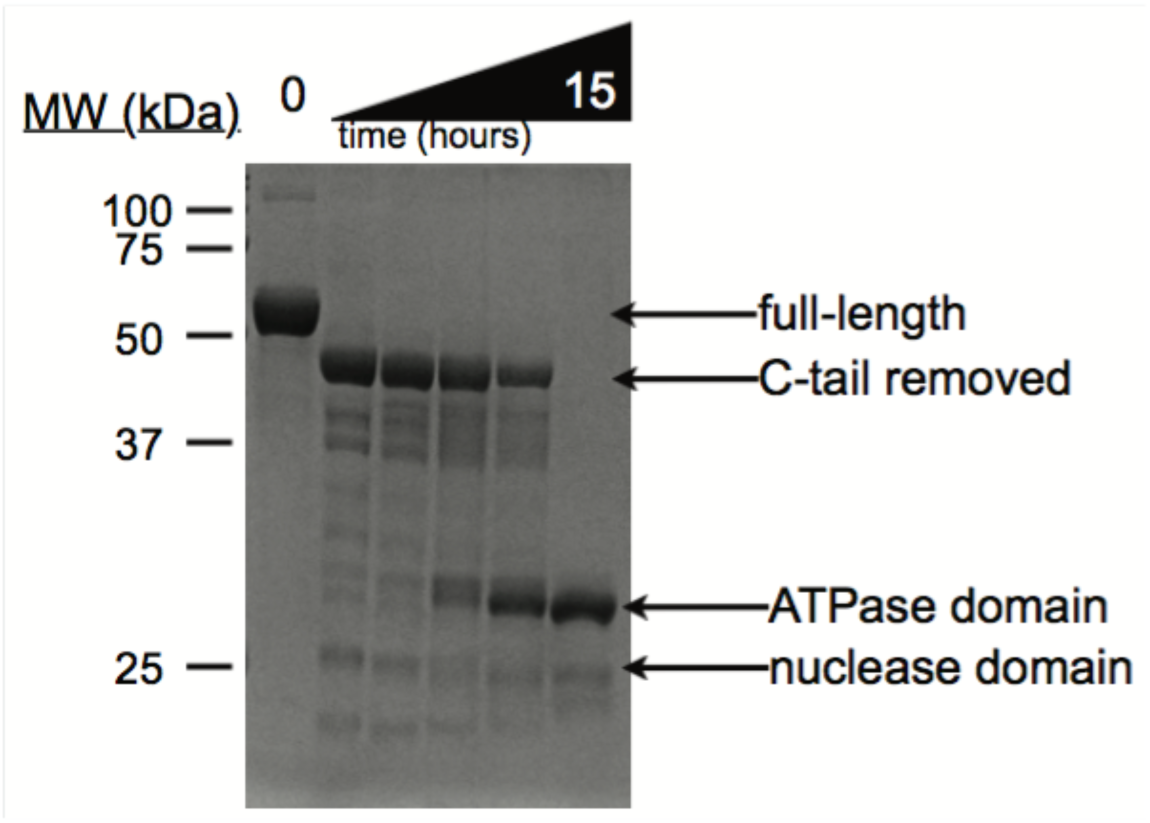
Limited proteolysis of TerL^P74-26^ rapidly cleaves the flexible C-terminal tail, including the disordered residues in our nuclease domain structure (Figure 5). Cleavage then occurs at the flexible linker between the ATPase and nuclease domains.

**Supplmental Table 1.**
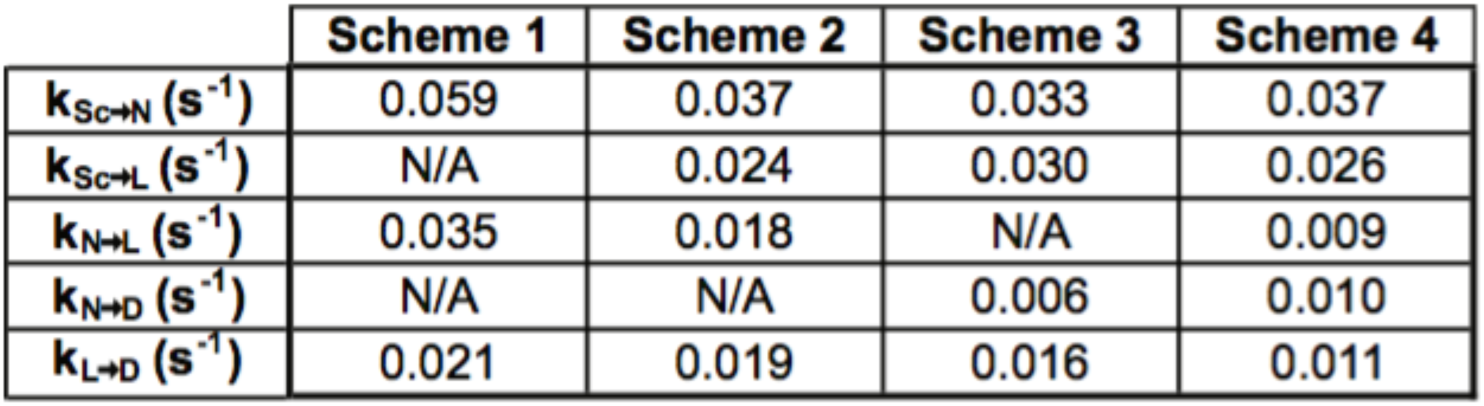
Rates obtained from kinetic modeling of nucleolysis data using Berkeley Madonna.

**Supplmental Table 2.** RMSD of the TerL^P74-26^-ND structure to similar structures.

| Phage/Virus | RMSD (Å) |
| --- | --- |
| T4 | 1.5 |
| Sf6 | 2.2 |
| P22 | 1.8 |
| SPP1 | 2.2 |
| CMV | 1.9 |
| HSV-1 | 2.1 |

**Supplmental Table 3.**
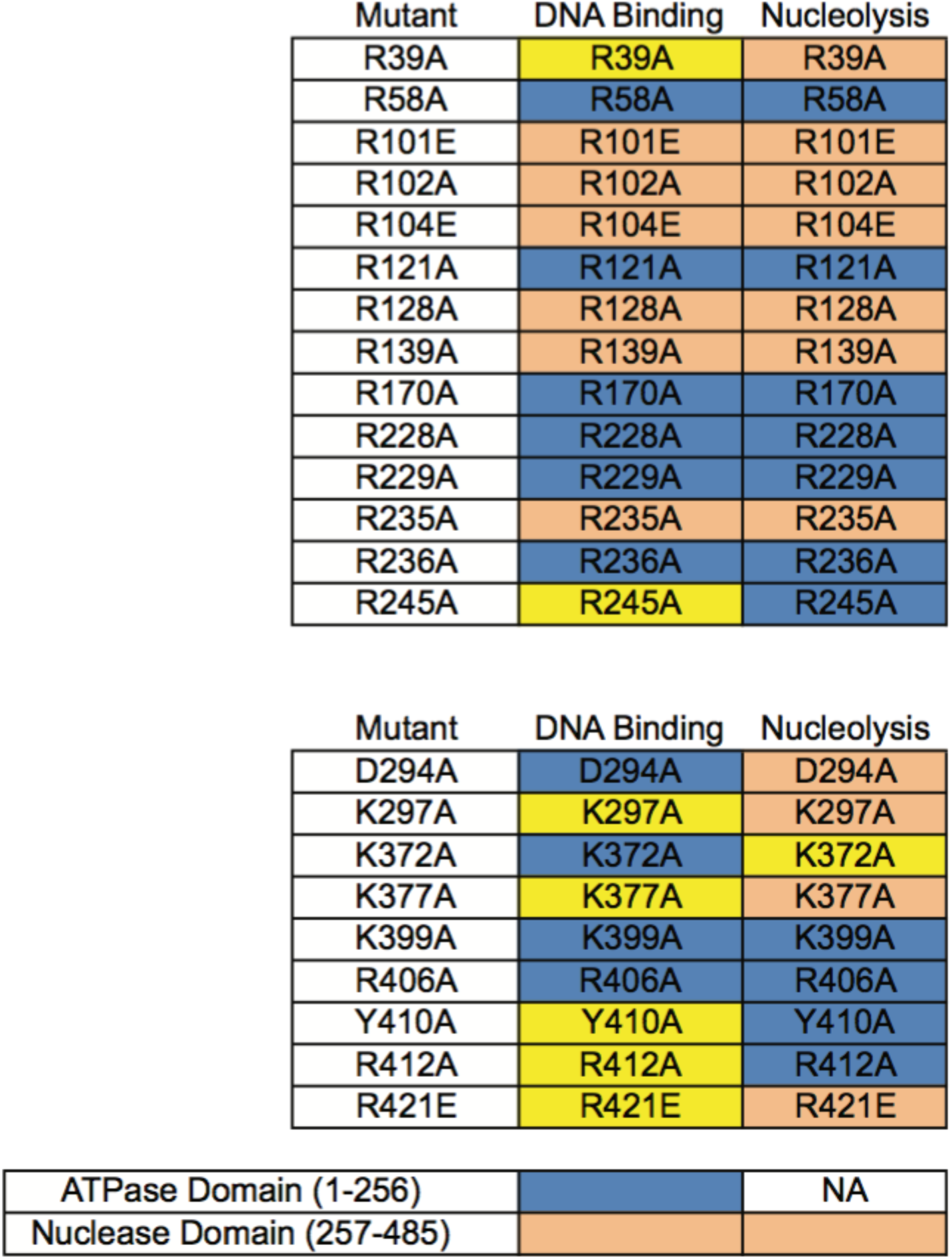
Effect of each mutation on DNA binding or cleavage. Variants with mild or no defect are colored blue. Variants with moderate or sever defects are colored yellow and orange respectively.

